# Impaired formation and updating of internal predictive models in a rat model of Fragile X Syndrome

**DOI:** 10.64898/2026.08.20.746021

**Authors:** D. Walker Gauthier, Ethan Hong, Noelle James, Cameron Killiam, Benjamin D. Auerbach

**Affiliations:** Department of Molecular and Integrative Physiology, School of Molecular and Cellular Biology, University of Illinois Urbana-Champaign, Urbana, Illinois, 61801, United States; Beckman Institute for Advanced Science & Technology, University of Illinois Urbana-Champaign, Urbana, Illinois, 61801, United States; Neuroscience Program, University of Illinois Urbana-Champaign, Urbana, Illinois, 61801, United States; Department of Neuroscience, Baylor College of Medicine, Houston, Texas, 77030, United States; National Eye Institute, National Institutes of Health, Bethesda, Maryland, 20892

**Keywords:** Auditory processing, predictive coding, oddball task, autism, deviant detection

## Abstract

Predictive coding frameworks propose that perception emerges from a continuous comparison of incoming sensory signals with internally generated predictions, with mismatches between the two computed as prediction errors. Disruptions to the balance between these top-down predictions and bottom-up sensory signals are theorized to contribute to sensory abnormalities in neuropsychiatric conditions like autism spectrum disorders. However, disambiguating bottom-up from top-down contributions to sensory processing remains a difficult challenge, particularly in animal models. Here we develop a probabilistic oddball detection task in which rats must track local statistics within a trial to detect a deviant stimulus, as well as global statistics across trials to anticipate when a deviant will occur. This design enables formation of experimentally specified internal models of deviant expectation that can be quantitatively derived from behavior and manipulated independently of local stimulus statistics. We used this task to characterize sensory predictive behavior in a *Fmr1* KO rat model of Fragile X Syndrome, the most common monogenic cause of autism. Male *Fmr1* KO rats detected deviant stimuli at wildtype levels but exhibited reduced anticipation of deviant occurrence based on cross-trial statistics and failed to adapt their behavior when these statistics changed. Computational modeling revealed that these behavioral deficits reflected an overweighting of immediate sensory evidence and temporal response pressure at the expense of learned predictive expectations, consistent with imprecise internal models skewed towards sensory immediacy. These findings provide evidence for disrupted predictive processing in Fragile X Syndrome, consistent with active inference accounts of autism, and highlight the utility of this probabilistic oddball task design for interrogating predictive coding and perceptual impairments in neuropsychiatric conditions.

## INTRODUCTION

We are constantly bombarded with a barrage of stimuli from the external world. To efficiently represent this overwhelming amount of information, our sensory systems must prioritize the encoding of certain inputs expected to provide meaningful outcomes. To achieve this, the brain is thought to build internal models of the world in order to identify and predict relationships between sensory objects, with these models being continuously updated in an experience-dependent manner (Friston, 2010). This predictive processing framework offers a unifying account of how sensory representations can be efficiently extracted from incomplete or noisy sensory input. Moreover, disruptions to predictive model formation or utilization have the potential to account for a wide range of sensory abnormalities observed across neuropsychiatric conditions (Qela et al., 2025). In autism spectrum disorders (ASD), for example, it has been proposed that atypical sensory processing, which is among the most prevalent and debilitating features of the condition (Sinclair et al., 2017; Thye et al., 2018), arises from a mis-weighting between top-down predictive models and bottom-up sensory signals (Pellicano and Burr, 2012; Lawson et al., 2014).

Fragile X syndrome (FXS) is a leading inherited cause of ASD and consistently presents with auditory processing issues (Rotschafer and Razak, 2014; McCullagh et al., 2020). We have previously demonstrated that a *Fmr1* KO rat model of FXS exhibits increased sound sensitivity and degraded feature encoding that is associated with enhanced cortical gain (Auerbach et al., 2021; Gauthier et al., 2025). When viewed from a predictive coding lens, these auditory phenotypes could be consistent with an overweighting of bottom-up sensory information relative to top-down expectations from internal models. However, disentangling bottom-up sensory processing from top-down predictive influences requires experimental paradigms that can manipulate internally generated expectations while providing quantitative behavioral readouts of sensory prediction. This is further compounded by the fact that predictive processing unfolds over multiple timescales and levels of abstractions (Friston, 2005; Kiebel et al., 2008), necessitating paradigms that can independently probe predictions at distinct levels of this hierarchy.

Oddball paradigms — in which a rare deviant stimulus is embedded within a sequence of standard stimuli — offer a naturalistic framework for probing predictive processing, as deviant-evoked responses are thought to reflect prediction error signals generated when incoming sensory input violates internal expectations (Nelken and Ulanovsky, 2007; Näätänen et al., 2007). Standard oddball designs, however, are unable to isolate lower-order statistical inferences, such as basic deviant detection driven by local stimulus probabilities, from higher-order, cross-trial inferences that reflect accumulated global regularities. While global-local oddball tasks have been developed to address this distinction (Bekinschtein et al., 2009), translating these approaches to provide quantitative behavioral readouts of predictive processing in rodent models remains a significant challenge.

To address this gap, we developed a probabilistic oddball paradigm, adapted from a human study (Tabas et al., 2020), that dissociates basic deviant detection within a given trial from deviant expectation based on broader statistical regularities across trials, enabling direct behavioral quantification of sensory expectation in rodents. Using this paradigm, we found that male *Fmr1* KO rats exhibited impaired deviant anticipation based on cross-trial statistics and failed to adapt their behavior when these statistics changed. Drift diffusion modeling revealed that these behavioral deficits reflected a bias towards immediate sensory evidence and temporal response pressure at the expense of accumulated prior expectations. Together, these results provide evidence that atypical auditory processing in FXS reflects a bottom-up overweighting of sensory evidence at the expense of top-down predictive expectations, consistent with a hypopriorism account of ASD, and establishes a tractable experimental framework for quantitatively characterizing predictive processing in rodent models of neuropsychiatric disorders.

## MATERIALS AND METHODS

### Subjects

Adult (>3 month old) male *Fmr1*^-/y^ (KO) rats on an outbred Long Evans (LE)background (LE-Fmr1^em2Mcwi^; Medical College of Wisconsin) and littermate wild-type (WT) controls were used for these studies. 13 WT and 10 *Fmr1* KO rats were used in total. Male rats were used because FXS occurs more frequently and in greater severity in males due to the X-linked nature of the disorder (Reiss and Hall, 2007). Rats lived in group-housed caging in a colony room maintained at 22□°C with a 12-h light–dark cycle. All subjects had free access to food and water except for those undergoing operant conditioning, when rats were food restricted and kept at approximately 90% of their free-feeding weight. Food-restricted animals had unrestricted access to water, except while participating in behavioral testing. Testing sessions lasted approximately 1 hour per day, with rats participating in one behavioral testing session per day, 6 days per week. All experiments were approved by the University of Illinois Urbana-Champaign Institutional Animal Care and Use Committee (IACUC) under protocol 23223, in accordance with NIH guidelines.

### Operant Conditioning

Rats were trained in a Go/No-Go operant conditioning paradigm using procedures similar to those described in our previous publications (Auerbach et al., 2021; Gauthier et al., 2025). Testing was carried out in operant conditioning chambers (Med-Associates, Inc., Model ENV-008-VP, St. Albans, VT) equipped with pellet dispensers (Med-Associates Model ENV-203 M, Fairfax, VT), illuminated nose-pokes with infrared sensor (Med-Associates, Inc., Model ENV-114BM, St. Albans, VT), and cue lights for signaling task status (Med-Associates, Inc., Model ENV-215M-LED, St. Albans, VT). All components were housed in single-walled sound-attenuating cubicles equipped with ventilation (Med-Associates, Inc., Model ENV-018MD, St. Albans, VT). Each behavioral setup was independently controlled by a dedicated microcontroller (Arduino DUE, Italy) that recorded signals from the behavioral devices, responded according to predefined protocols, and communicated with a host computer to transmit control signals and data.

The entire system was controlled by custom software written in MATLAB (MathWorks, R2022, USA), which handled system control, stimulus generation, and data collection and visualization. Sound stimuli (192 kHz sampling rate) were generated and delivered using a multi-channel sound card (RME MADIface Pro, Germany), multi-channel digital-to-analog converter (RME M-32 DA Pro, Germany), power amplifiers (Behringer NX1000, Germany), and speakers (Fostex FT17H Horn Super Tweeter, Japan) positioned approximately 30 cm above the animal’s head. Stimuli were calibrated using a microphone preamplifier (Larson Davis, model 2221, Depew, NY) equipped with a ½” microphone (Larson Davis, model 2520, Depew, NY) at a location where the animal’s head would be during a trial.

Rats were first trained to detect tone bursts (8 kHz, 50 ms duration, 5 ms rise/fall time, cosine gated) (**Supplemental Fig 1A**). A rat began a trial by placing its nose in a nose-poke hole, which initiated a variable waiting interval ranging from 1 to 4□s. During the waiting interval, the rat had to maintain its position in the nose-poke hole until it heard a tone burst or the trial was aborted. In the Go condition, the target stimulus was the tone burst. If the rat detected this signal, it removed its nose from the nose-poke hole resulting in a food reward (45□mg dustless rodent grain pellets, Bio-Serv); a hit was recorded if the rat correctly responded to the tone within 2□s. A miss was recorded if the rat failed to remove its nose from the nose-poke within the 2□s response interval. No reinforcement was given for a miss. Approximately 30% of all trials were catch trials, where tone bursts were not presented. This constituted the No-Go part of the procedure. If the rat removed its nose during a catch trial, a false alarm (FA) was recorded and the rat received a 4-8□s timeout, during which the house light was turned off and the rat could not initiate another trial. However, if the rat continued to maintain nose-poke position during these catch trials, a correct rejection (CR) was recorded. No reinforcement was given for a correct rejection.

After initial detection training using a 70 dB SPL Go stimulus (training criteria: > 240 trials, > 90% hit rate, <20% FA rate over 3 consecutive days), the animals started discrimination training, where a 60 dB SPL broadband noise stimulus (BBN, 1-65 kHz, 50 ms duration, 5 ms rise/fall time, cosine gated) was introduced for No-Go catch trials (**Supplemental Fig 1B**). After meeting performance criteria (training criteria: > 240 trials, > 90% hit rate, <20% FA rate over 3 consecutive days), animals were next trained to report the detection of a Go tone embedded within a series of 6 No-Go BBN stimuli (**Supplemental Fig 1C**). This constituted the oddball or deviant detection phase of training.

Oddball trials were self-initiated via nose-poke. Following a 1-2 s variable delay period, a train of seven stimuli were presented, consisting of six BBN No-Go stimuli (standards) and one 8 kHz Go tone (deviant) with an ISI of 700 ms. A deviant tone was present once in every trial, occurring with equal probability in positions 4, 5, or 6 within the train of stimuli (**Fig 1A**). If the animal responded within the 700 ms response window immediately following the deviant presentation, this was reported as Hit and the animal received a food reward. If the animal responded to any standard BBN stimulus before deviant presentation, this was counted as a FA, and the animal was punished with an 8s timeout. If the animal failed to respond within the 700 ms response window immediately following the deviant presentation, this was reported as a miss. No reinforcement was given for misses.

**Figure 1.**
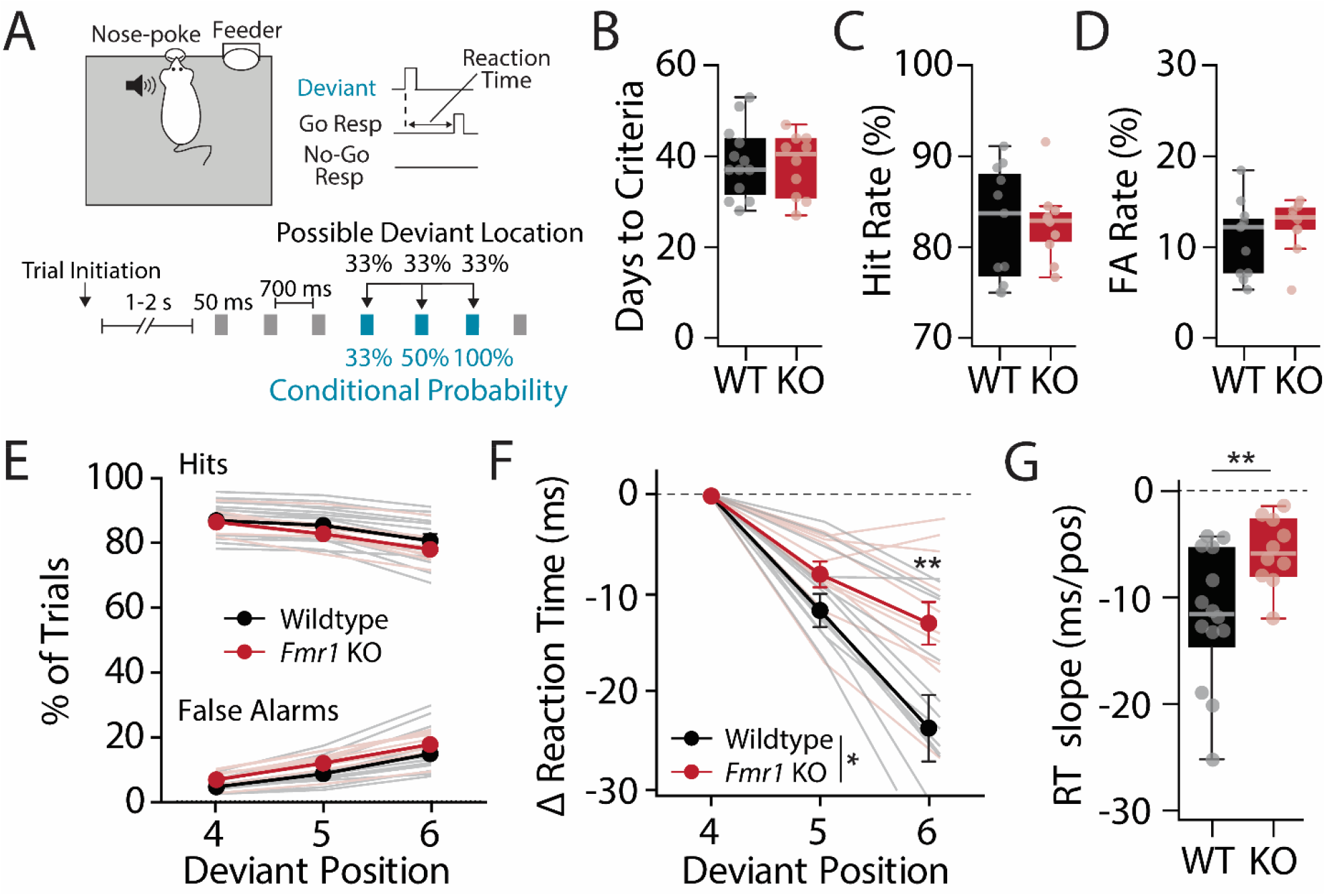
*Fmr1* KO rats show intact deviant detection but impaired deviant expectation in a probabilistic oddball task. **(A)** Experimental design. Male wildtype (n = 13) and *Fmr1* KO (n = 10) rats were operantly trained to detect a deviant tone stimulus (blue) embedded within a series of broadband standards (gray). Self-initiated trials consisted of seven consecutive stimuli of 50 ms duration with an interstimulus interval of 700 ms. In the baseline phase, a single deviant stimulus was guaranteed to occur on every trial at either position 4, 5, or 6 with equal probability. In this manner, deviant occurrence becomes increasingly predictable at later positions within a trial (see conditional probability). **(B)** Box plot showing total training time across all phases of task acquisition (detection, discrimination, oddball) for wildtype (black) and *Fmr1* KO (red) rats. **(C)** Hit rates and **(D)** false alarm (FA) rates for fully trained WT and *Fmr1* KO rats. **(E)** Hit and FA rates for WT and *Fmr1* KO rats as a function of deviant position. **(F)** Changes in response latency as a function deviant position (normalized to position 4 reaction time). **(G)** Box plot of resultant fit linear slope values to reaction time x position curves. *p < 0.05, **p < 0.01, ***p < 0.001, ns = not significant.

Once criteria (>200 trials, >75% hit rate, < 20% FA rate over 3 consecutive days) was reached on this training version of the oddball task, the frequency of Go tone was rotated between 4, 8, 16, and 32 kHz for each behavioral session in pseudorandomized fashion. Upon successful criteria across all 4 frequencies, oddball training was considered to be completed, and baseline phase data collection proceeded with the same randomized rotation of deviant frequency. A total of 13 WT and 10 *Fmr1* KO rats were tested on the baseline phase of the task across three cohorts (**Supplemental Fig 2A**). Following the baseline phase, cohort 1 (n = 4 WT, 3 *Fmr1* KO rats) was introduced to the probe phase, where 2 sessions per week contained rare probe trials consisting of 7 standard BBN stimuli and no deviant tone stimulus (∼1% of total trials). These probe trials were unreinforced, i.e. incorrect responses to BBN standards (FAs) were not punished. The probe phase was conducted until at least 40 FAs were recorded for each animal. Cohort 2 (n = 5 WT, 3 *Fmr1* KO) did not undergo probe phase but was instead introduced to the catch phase following baseline testing. During the catch phase, the proportion of trials containing no deviant stimulus was increased to 40% of the overall trials, and unlike in the probe phase, animals were punished with a timeout for incorrect responses to BBN standards (FAs). Catch sessions were presented daily for at least 7 consecutive behavioral sessions. Cohort 3 (n = 4 WT, 4 *Fmr1* KO) underwent all three phases of testing (baseline → probe → catch). Thus, a total of 8 WT and 7 *Fmr1* KO rats were subjected to both baseline and probe phases of the task, while a total of 9 WT and 7 *Fmr1* KO rats were subjected to both baseline and catch phases of the task. Importantly, similar behavioral performance was observed across all three cohorts (**Supplemental Fig 2B-D**).

### Computational Modeling

We developed a drift-diffusion model (DDM) to estimate decision-making parameters that could account for genotype differences in behavioral output (see **Fig. 4A**). The model incorporates three contributions to responding: stimulus-dependent evidence, an expectation component determined by the probability of deviant occurrence, and a general pressure to respond that increases with elapsed time. These components jointly determine the accumulation of evidence toward a response threshold, allowing the model to predict both response timing and the probability of withholding a response. Parameters were optimized separately for each animal using baseline and catch-phase data. Animals were included if they had completed both phases, yielding a modeling cohort of 9 WT and 7 *Fmr1* KO rats.

#### Evidence Accumulation and Response Generation

Within each trial, a decision variable *X_t_* accumulated according to:

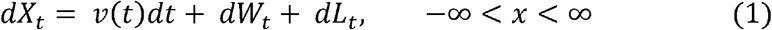

where *v(t)* represents the time-dependent drift, *W_t_* is a Wiener process with noise scale fixed to one, and *L_t_* imposes a reflecting boundary at zero. A decision occurred when accumulated evidence first reached an absorbing threshold *a*. The predicted response time was defined as:

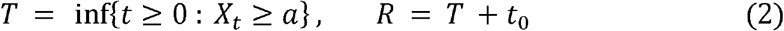

where *t*_0_ represents a constant nondecision delay. Responses were classified as hit, miss, false alarm, or correct rejection by the same criterion used in the behavioral task (see Operant Conditioning Methods). Nonresponses (misses and correction rejections) contributed their survival probability to the likelihood and were not assigned an observed reaction time. Prior to presentation of a deviant, drift was determined by:

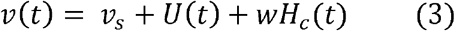

where *v_s_* is the standard-stimulus drift, *U(t)* is elapsed-time response pressure, and *w* determines the contribution of the expectation signal *H_c_(t)* under the task phase parameter *c*, which denotes either baseline or catch phase. Response pressure increased linearly until the final possible deviant onset and then relaxed if no deviant occurred:

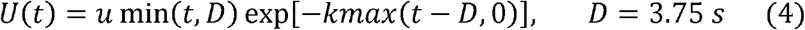

Here, *u* controls the increase in response pressure and *k* controls its subsequent decline. When *k=0*, response pressure remains constant after the final opportunity. When *u=0*, the decay parameter has no effect on model predictions.

Following an actual deviant at position *j*, drift became:

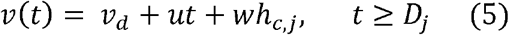

where *v_d_* is the deviant-stimulus drift and *h_c,j_* is the expectation value associated with that position. Thus, following a deviant, the original rising time-pressure term continued through the response window. The derived quantity *v_d_-v_s_* describes the modeled change in drift associated with deviant presentation, which we term deviant salience.

#### Representation of Deviant Expectation

The expectation component was constructed from the conditional probability of deviant occurrence given that no earlier deviant had appeared:

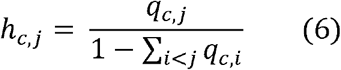

where *q_c,j_* is the scheduled probability of a deviant at position *j*. Deviant expectation was calculated from the stimulus probabilities specified for each task phase (see Operant Conditioning Methods), conditional on no deviant having appeared earlier in the trial. These conditional probabilities defined the deviant expectation signal, whose contribution to drift was scaled by the fitted expectation-weight parameter, *w*. Phase-dependent predictions arise from changes in the conditional probability signal (Hc(t)), which is derived directly from the task design and therefore differs between baseline and catch phases, reflecting the conditional probability distributions of deviant occurrence for each phase. The expectation weight thus reflects how strongly an animal integrates position-dependent changes in deviant probability into its decision process, regardless of which phase those probabilities are drawn from, such that animals with higher expectation weights show greater sensitivity to phase-dependent changes in task statistics.

The expectation signal reflected the conditional probability of the next possible deviant during the 0.75 s before its onset. It was zero outside these anticipation periods unless a deviant occurred, in which case its value was maintained through the response window. The nondecision delay, t□, was then added to predicted decision times to obtain response times.

#### Model Fitting and Validation

Trial identifiers, stimulus timing, response labels, and task schedules were audited before fitting. The resulting analysis included 736,190 trials. For each animal, model training was performed on the first 93% of baseline sessions as well as the first half of the catch phase sessions. Seven parameters were fitted per animal: standard drift, deviant drift, response threshold, nondecision delay, expectation weight, time-pressure slope, and urgency decay. Model parameters were shared across baseline and catch phases within each animal; phase-dependent predictions arose from the corresponding taskschedules. Parameters were optimized by minimizing the negative log-likelihood of responses in 10-ms bins and nonresponses at the trial deadline. Optimization used differential evolution with multiple random seeds and search budgets, followed by bounded local refinement. The parameter vector yielding the highest training likelihood was retained.

The DDM was evaluated using simulated parameter recovery, optimization stability, numerical accuracy, and prediction of later-session behavior (**Supplemental Fig 4**). Across 20 simulated datasets for each animal, parameters recovered close to their generating values with an average correlation of r = 0.8892 across parameters. The decay parameter was poorly recovered for one animal, though the value was near zero for this subject, making recovery inherently difficult. The model was stable across initialization conditions, and longer searches produced equivalent or identical likelihoods, confirming optimization stability. All 48 numerical resolution checks also passed. Together these results support the validity of the model fitting procedure and the reliability of the parameter estimates.

### Statistics

Statistical analyses were performed in Python (3.10.0). Normality was assessed using the Shapiro-Wilk test prior to selecting parametric or non-parametric alternatives. For single pairwise comparisons, Welch’s t-tests (unequal variance) were used when data were normally distributed; Mann-Whitney U tests were used otherwise. Where one-sided tests were applied, directional hypotheses were specified a priori based on theoretical predictions. For multiple pairwise comparisons, family-wise error was controlled using either Bonferroni correction or Benjamini-Hochberg false discovery rate (FDR) correction, as specified per analysis. Repeated measures ANOVAs were used for within-subject comparisons across timepoints or positions, with Greenhouse-Geisser correction applied when sphericity was violated (ε < 1). Post-hoc paired t-tests with Holm-Bonferroni correction or Wilcoxon signed-rank tests were used following significant omnibus tests, as appropriate. For computational model parameters, genotype comparisons used one fitted parameter vector per animal. Differences in model parameters were evaluated using Mann-Whitney U test. All statistical values are reported as means ± standard error of the mean (SEM), unless otherwise stated. Boxplots display the median, 25th–75th percentiles (box), full range (whiskers), and outliers (below Q1 − 1.5×IQR or above Q3 + 1.5×IQR). Boxplot dots correspond to individual animal level data or parameter fits. Statistical details for all figure panels are provided in **Supplemental Table 1**.

## RESULTS

### Probabilistic oddball task produces behavioral readout of sensory expectation

To examine predictive processing in rodents, we developed an operant-based oddball detection paradigm where deviant expectation can be manipulated independently of local stimulus statistics (Tabas et al., 2020). In this task, subjects self-initiate trials consisting of a sequence of 7 sounds — 6 standard broadband noise bursts and 1 deviant oddball tone (4, 8, 16, or 32 kHz)— and report detection of the deviant in order to receive a food reward (**Fig 1A**). In the baseline phase, every trial contained one deviant stimulus that occurred at position 4, 5, or 6 within the sequence at equal probabilities. In this manner, deviant occurrence becomes increasingly predictable as the sequence progresses, with the conditional probability of deviant occurrence rising from 33% to 100% from position 4 to 6 within a given trial (**Fig 1A**). This design allowed us to independently assess two distinct components of predictive processing: (1) the ability to detect a deviant stimulus based on local trial statistics (deviant detection); and (2) the ability to track global regularities in deviant occurrence across trials to anticipate when a deviant will occur (deviant expectation). Critically, deviant expectation can be manipulated independently of stimulus features and local statistics in this task, allowing us to dissociate deficits in high-order predictive model formation from alterations in basic deviant detection (Tabas et al., 2020).

Male *Fmr1* KO (n = 10) and littermate WT (n = 13) rats learned this deviant detection task at comparable rates (**Fig 1B**; Mann-Whitney U test: U = 64.5, p = 1.000). There was no difference in task acquisition rate or days to criteria between WT and *Fmr1* KO rats for any phase of training (**Supplemental Fig 1**). These results are consistent with our previous findings showing normal acquisition of operant detection and discrimination tasks in *Fmr1* KO rats (Auerbach et al., 2021; Gauthier et al., 2025). In regard to deviant detection abilities, there was no significant genotype difference for fully trained animals, with WT and *Fmr1* KO rats exhibiting comparable hit rates (**Fig 1C**; Welch’s t-test: t = 1.279, p = 0.215), even when accounting for outliers (Welch’s t-test: t = 1.982, p = 0.063). There was a trend towards higher false alarm (FA) rates in *Fmr1* KO rats (**Fig 1D**; Mann-Whitney U test: U = 31, p = 0.060), which reached statistical significance upon outlier analysis (Mann-Whitney U test: U = 26, *p = 0.044). Notably, both genotypes maintained task performance at above criteria levels (hit rates > 75% and FA rates < 20%) for all deviant frequencies and all task phases. Additionally, we found no differences in trial initiation rate or attempts per trial between WT and *Fmr1* KO animals, measures that provide implicit indices of task motivation and impulsivity (**Supplemental Fig 1**). Together, these results indicate that WT and *Fmr1* KO subjects are both adept at detecting deviant stimuli in this task, allowing for examination of deviant expectation.

If our subjects can track global regularities in deviant occurrence, we reasoned that task performance should systematically vary with deviant position, as deviant predictability increases across the sequence (**Fig 1A**). Hit and FA rates did indeed vary as a function of deviant position (mixed ANOVA, position: Hit rate: F(2,42) = 95.85, ****p < 0.0001; FA rate: F(2,50) = 115.99, ****p < 0.0001), with hit rates decreasing and FA rates increasing for later occurring (and thus more predictable) deviants. This result was counterintuitive, as performance appeared to get worse for more predictable deviant stimuli. However, examination of reaction times as a function of deviant position clarified this finding (**Fig 1F**). To examine how reaction times changed as a function of deviant position (and thus predictability), we examined the difference in reaction times to deviants occurring at position 5 and 6 relative to position 4 (**Fig 1F**).

In WT rats, reaction times became progressively faster for later occurring deviants, as indicated by the steep slope of their reaction time curves (**Fig 1G**). This decrease in reaction time and increase in false alarm rate seen at later positions may thus reflect anticipatory responding, with animals expecting the deviant so strongly at position 6 that they respond in preparation for its arrival. Intriguingly, *Fmr1* KO rats did not modulate their reaction times as a function of deviant position to the same degree as WT animals (**Fig 1F**; mixed ANOVA: genotype F(2,40) = 6.64, *p = 0.012; genotype x position F(2,42) = 4.673, *p = 0.015), with a significantly shallower slope to their reaction time curves (**Fig 1G**; Welch’s t-test: t =−3.12, **p = 0.006). Notably, *Fmr1* KO animals exhibited significantly faster absolute reaction times compared to WT littermates (**Supplemental Fig 3A;** mixed ANOVA: genotype F(1,21) = 5.739, *p = 0.026), consistent with our previous findings (Auerbach et al., 2021). However, there was no significant correlation between overall reaction time and position-dependent changes to reaction time in either genotype (WT, Pearson r = 0.117, p = 0.747; KO: Pearson r = 0.038, p = 0.903), indicating that the shallower reaction time curves seen in *Fmr1* KO rats are unlikely to be due to a “floor effect” (**Supplemental Fig 3B**). Indeed, when analysis was restricted to subjects whose overall reaction time fell within ±1 SD of the combined group mean (thereby removing genotype differences in absolute reaction time), a significant genotype difference in position-dependent reaction time changes remained (**Supplemental Fig 3C-E**). These results suggest that WT animals more strongly anticipate deviant occurrence and prepare motor responses compared to *Fmr1* KO rats.

### *Fmr1* KO rats exhibit deficits in sensory expectation and updating

The position-dependent reaction time changes described above could indicate that animals anticipate deviant occurrence based on learned internal models, with this capacity diminished in *Fmr1* KO rats. To further examine whether animals were utilizing top-down predictive models in this task, we introduced a probe phase to a subset of trained animals (n = 8 WT, 7 *Fmr1* KO rats) in which rare probe trials containing no deviant stimulus were embedded within behavioral sessions (**Fig 2**). These probe trials constituted only 1% of total trials and were unreinforced, allowing us to covertly observe anticipatory behavior without altering the conditional probability structure of the task (**Fig 2A**). Hit rates modestly improving during the probe phase (mixed ANOVA: phase, F(1,13) = 5.026, *p = 0.043); however, no effect of genotype (mixed ANOVA: genotype, F(1,13) = 2.664, p = 0.127) or genotype x phase interaction (mixed ANOVA: genotype x phase, F = 2.567, p = 0.133) were observed (**Fig 2B**). The introduction of probe trials did not alter overall FA rates (**Fig 2C**; mixed ANOVA: phase, F(1,13) = 0.035, p = 0.854; genotype, F(1,13) = 3.358, p = 0.090; genotype x phase interaction, F(1,13) = 1.018, p = 0.331). Importantly, FA rate on probe trials was also well within performance criteria (<20%) for both genotypes and comparable to FA rates on normal deviant-containing trials (**Fig 2D**). Finally, *Fmr1* KO rats continued to exhibit a shallower reaction time curve relative to WT rats in the probe phase (**Fig 2E**; genotype × position mixed ANOVA: genotype, F(1,13) = 6.77, *p = 0.022; genotype x position, F(2,26) = 7.826, **p = 0.002), with a significantly steeper reaction time slope in WT compared to *Fmr1* KO animals across phases (**Fig 2F**; mixed ANOVA: genotype, F(1,13) = 11.94, **p = 0.004) but no difference in slope within genotype across phase (mixed ANOVA: phase, F(1,13) = 0.204, p = 0.659; genotype x phase, F(1,13) = 0.660, p = 0.431). These results confirm that introduction of probe trials does not alter basic task performance, thus providing an opportunity to read out anticipatory internal models of deviant occurrence.

**Figure 2.**
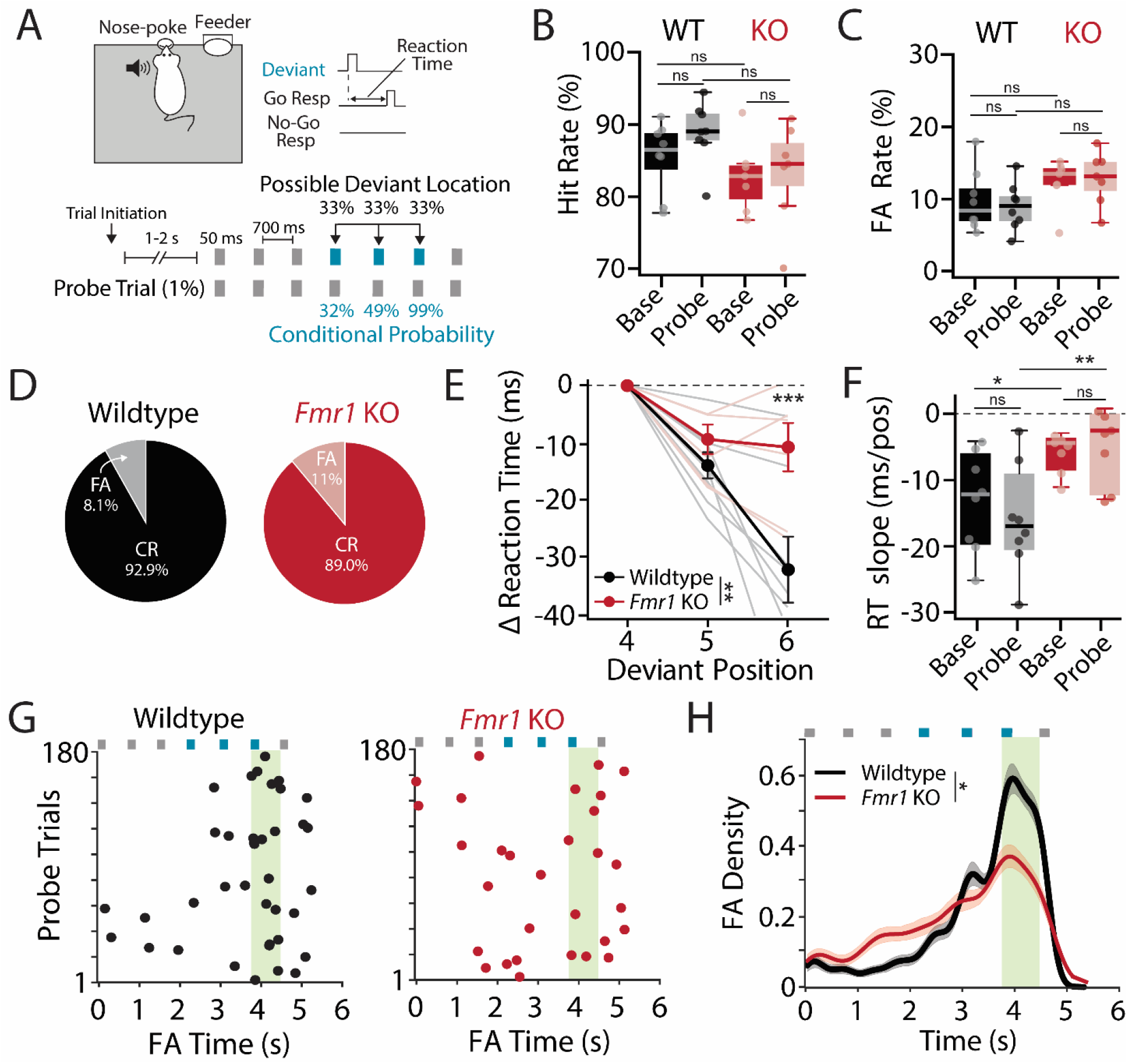
Covert probe trials reveal a precise internal model of deviant timing in WT rats that is degraded in *Fmr1* KO rats. **(A)** Experimental design for probe phase (n = 8 WT, 7 *Fmr1* KO rats). During the probe phase, there is a 1% chance of an unreinforced probe trial that does not contain a deviant stimulus. **(B)** Hit rates compared between baseline and probe phases for WT (black/gray) and *Fmr1* KO (red/pink) rats. **(C)** False alarm rate compared between baseline and probe phases for WT and *Fmr1* KO rats. **(D)** Pie charts illustrating false alarm (FA) and correct rejection (CR) rate for WT and *Fmr1* KO rats on probe trials specifically. **(E)** Deviant response latency normalized to position 4 reaction time during the probe phase. **(F)** Comparison of normalized reaction time x position slope between baseline and probe phase for WT and *Fmr1* KO rats. **(G)** Raster plots showing temporal distribution of FAs on probe trials from an example WT (left) and *Fmr1* KO (right) rat. Each row represents an individual probe trial, dots reflect a FA at a given time on a given probe trial. **(G)** Population distribution of FA timing for WT and *Fmr1* KO animals on probe trials. Green bar represents position 6 response window. *p < 0.05, **p < 0.01, ***p < 0.001, ns = not significant.

Because no deviant tone is presented during probe trials, the temporal distribution of false alarms on these trials may reflect the animal’s internal expectation of when a deviant should occur. Figure 2G shows raster plots of false alarm (FA) timing during probe trials for representative WT and *Fmr1* KO rats. Population distributions of FA time for each genotype are quantified in Figure 2H. WT rats exhibited a precise FA distribution that ramped up across trial time, peaked sharply at the onset of position 6 (when deviant occurrence is most likely), and then tapered off thereafter (**Fig 2H**). Notably, FA rate declined after position 6 rather than continuing to increase through position 7, indicating that animals had encoded position 6 as the most expected deviant location rather than exhibiting a general pressure to respond. This pattern is indicative of a precise internal model of deviant expectation. In contrast, *Fmr1* KO rats exhibited a markedly less precise false alarm distribution during probe trials compared to WT animals (Cluster Permutation KS-test: D = 0.192, *p = 0.028), characterized by a diminished peak and broader spread of FAs across stimulus positions (**Fig 2H**). These results suggest that WT rats learn to anticipate deviant occurrence based on global regularities across trials, while *Fmr1* KO animals exhibit a comparatively uniform FA distribution on probe trials, consistent with a less precise internal model of deviant position.

Having established that WT animals can anticipate deviant occurrence based on learned global statistics, and that this anticipation is blunted in *Fmr1* KO rats, we next asked whether animals could update their internal models in response to changes in global task statistics. To accomplish this, we introduced a catch phase to WT (n = 9) and *Fmr1* KO (n = 7) rats (**Fig 3**). During the catch phase, the proportion of trials containing no deviant stimulus was increased to 40% of the total trials and, unlike in the probe phase, animals were punished for FAs on these no-deviant catch trials. The introduction of catch trials thereby flattened the difference in conditional probability across deviant location (**Fig 3A**). Deviant occurrence was thus no longer predictable during the catch phase, rendering previously formed internal models of deviant expectation invalid and providing a direct test of model updating, a core feature of predictive processing frameworks (Keller and Mrsic-Flogel, 2018).

**Figure. 3.**
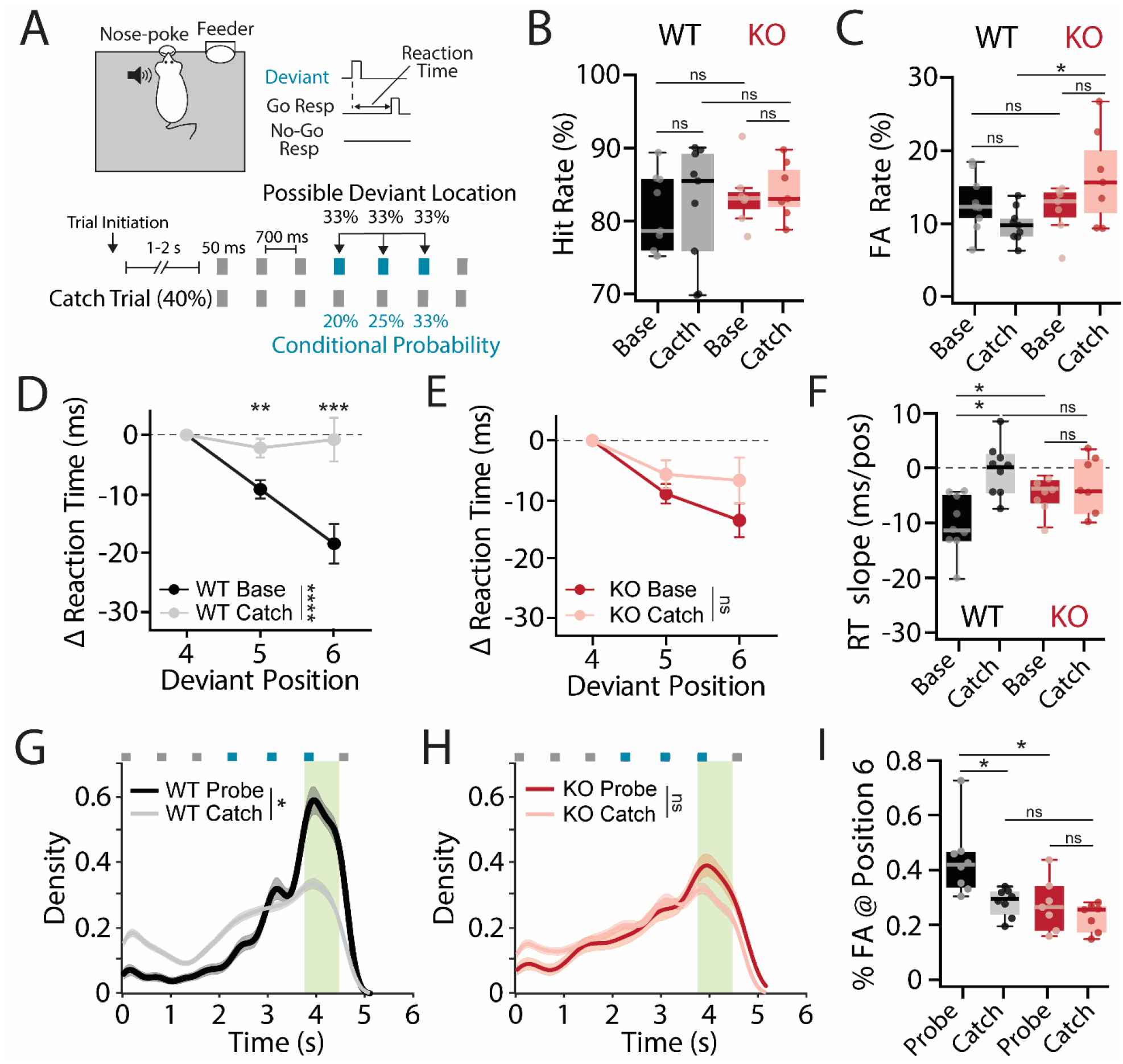
Catch phase change to cross-trials statistics produces internal model updates in WT but not *Fmr1* KO rats. **(A)** Experimental design for catch phase (n = 9 WT, 7 *Fmr1* KO rats). During the catch phase there is a 40% chance that a trial does not contain a deviant, compressing the change in conditional probability of deviant occurrence as a function of stimulus position. Response to any standard stimulus on these catch trials was punished with an 8s timeout. **(B)** Hit rates compared between baseline and catch phases for WT (black/gray) and *Fmr1* KO (red/pink) rats. **(C)** False alarm (FA) rates compared between baseline and catch phases for WT and *Fmr1* KO rats. **(D)** Deviant response latency normalized to position 4 reaction time for baseline and catch phase for WT rats (n = 9). **(E)** Deviant response latency normalized to position 4 reaction time for baseline and catch phase for *Fmr1* KO rats (n = 7). **(F)** Comparison of normalized reaction time x position slope between baseline and catch phase for WT (n = 9) and *Fmr1* KO (n = 7) rats. **(G)** Temporal distribution of false alarms on probe (black, n = 8) and catch (gray, n = 9) trials for WT rats. Green bar represents position 6 response window. **(H)** Temporal distribution of false alarms on probe (red, n = 7) and catch (pink, n = 7) trials for *Fmr1* KO rats. Green bar represents position 6 response window. **(I)** Comparison of the percent of FAs on probe (black/red) or catch (gray/pink) trials occurring within the position 6 response window in WT (probe: n = 8; catch n = 9) and *Fmr1* KO (probe: n = 7; catch: n = 7) animals. *p < 0.05, **p < 0.01, ***p < 0.001, ****p<0.0001, ns = not significant.

Introduction of reinforced catch trials did not affect hit rates in either genotype (**Fig 3B**; mixed ANOVA: phase, F(1,14) = 0.557, p = 0.468; genotype, F(1,14) = 0.8442, p = 0.374; phase x genotype, F(1,14) = 0.023, p = 0.881). Moving to the catch phase also did not significantly alter FA rates in either genotype, though it should be noted that FA rates were significantly higher in *Fmr1* KO compared to WT rats during the catch phase (**Fig 3C**; ANOVA: phase, F(1,14) = 0.509, p = 0.487; genotype, F(1,14) = 2.366, p = 0.146; phase x genotype, F(1,14) = 6.315, *p = 0.018). These results indicate that overall task performance was relatively preserved during catch phase.

We next determined how introduction of catch trials affected deviant expectation. In WT animals, reaction time curves were strikingly flatter compared to baseline phase, indicative of less anticipatory responding (**Fig 3D**; RM ANOVA: phase, F(1,8) = 34.81, ****p < 0.0001; phase x position, F(1,8) = 31.87, ***p = 0.0002). Intriguingly, reaction time curves did not significantly flatten in the catch phase for *Fmr1* KO rats (**Fig 3E**; RM ANOVA: phase, F(1,6) = 3.118, p = 0.066; phase x position, F(1,6) = 1.12, p = 0.331). Indeed, while reaction time slope was significantly reduced in WT animals between baseline and catch phases, no significant change was observed in *Fmr1* KO animals (mixed ANOVA: phase, F(1,14) = 15.77, **p = 0.001; genotype, F(1,14) = 0.237, p = 0.634; genotype x phase, F(1,14) = 6.873, *p = 0.020). The distribution of FAs on catch trials also became more evenly spread across stimulus positions in WT animals compared to the precise peak seen on probe trials, potentially reflecting adaptive updating of internal models when task statistics changed (**Fig3G**; Cluster Permutation KS-test: D = 0.266, *p = 0.032), while the FA distribution in *Fmr1* KO animals did not change between probe and catch trials either (**Fig 3H**; Cluster Permutation KS-test: D = 0.1189, p = 0.1238). To examine genotype-phase interactions in FA distribution, the percent of FAs occurring within the position 6 response window on catch trials was compared to the percent of FAs occurring within the position 6 response window on probe trials (**Fig 3I**). There was a significant main effect of genotype and phase on this measure (two-way ANOVA: phase, F(1, 27) = 8.79, p = 0.006; genotype, F(1, 27) = 10.93, p = 0.003), with the percentage of FAs occurring during position 6 significantly reduced in WT animals for catch trials compared to probe trials, while no significant change was detected in *Fmr1* KO animals across phase. However, the genotype × phase interaction was not significant (two-way ANOVA genotype × phase; F(1, 27) = 2.678, p = 0.114). Taken together, these results suggest that internal models of deviant occurrence can be manipulated via changes in global task statistics, with WT animals adapting their behavior to match the new conditional probability distribution while these changes were less pronounced in *Fmr1* KO animals. Thus, *Fmr1* KO rats not only exhibit less precise internal models of deviant expectation but also appear less able to update these models when global task statistics change.

### *Fmr1* KO rats exhibit a decision-making profile consistent with sensory immediacy

Our behavioral results are suggestive of deficits in the formation and flexible updating of predictive representations in a rodent model of FXS, consistent with theoretical accounts proposing that ASD is characterized by an over-reliance on incoming sensory evidence at the expense of top-down predictive models (Pellicano and Burr, 2012; Lawson et al., 2014). To more directly examine the internal parameters that could be driving genotypic differences in behavioral output, we developed a drift diffusion model (DDM) of the decision-making process underlying task performance (**Fig 4**, see Methods). DDMs are a well-validated framework for decomposing response time distributions and response probabilities into interpretable latent parameters reflecting distinct aspects of the decision process, including the rate of evidence accumulation (drift rate), the amount of evidence required before committing to a response (threshold), and pre-decisional sensory and motor processing delays (non-decision time) (Ratcliff, 1978; Gomez et al., 2007).

**Figure. 4.**
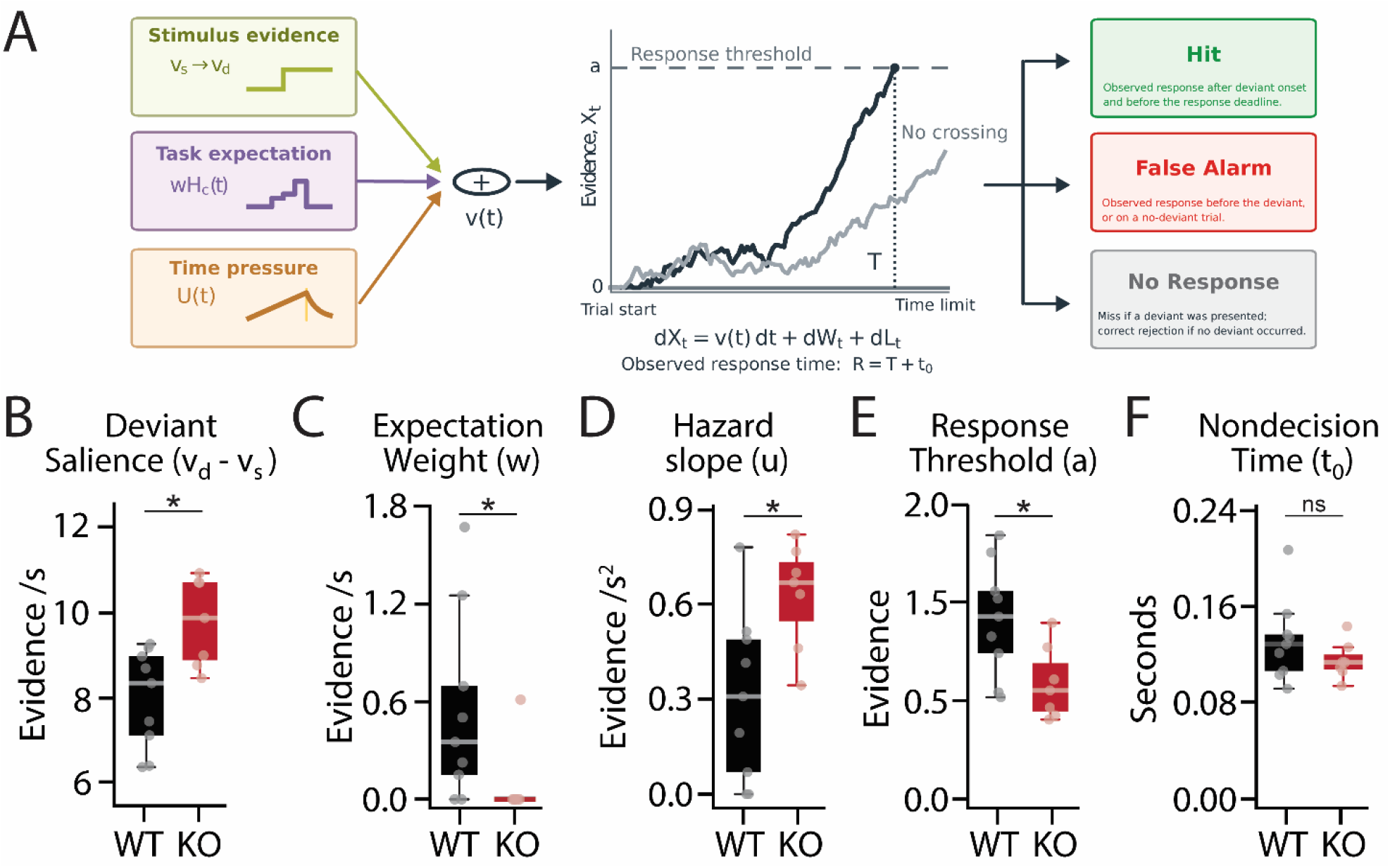
Characterizing perceptual decision making with a drift diffusion model. **(A)** Drift diffusion model schematic. The model decomposes evidence accumulation into three separable components: stimulus evidence, reflecting separately fitted drift rates for standard (v_s_) and deviant (v_d_) stimuli; deviant expectation weighting (wH_c_(t)), reflecting the contribution of learned predictive expectations to evidence accumulation; and temporal hazard (U(t)), reflecting time-dependent within-trial response pressure. A response is generated when accumulated evidence reaches threshold (a), with non-decision time (t□) representing pre-decisional sensory and motor processing delays. (**B-F)** Parameter fits for WT (black) and *Fmr1* KO (red) rats **(B)** Deviant-to-standard drift difference (v_d_ - v_s_). Higher values indicate an increase in evidence accumulation upon deviant presentation, reflecting perceptual salience of deviant. **(C)** Deviant expectation weighting. Higher values indicate that including conditional probabilities of deviant occurrence improve model fit, while values near zero indicate that behavior was better explained without an expectation component. **(D)** Hazard slope. Higher values indicate that a linearly increasing urgency signal accounts for a subject’s tendency to respond at later stimulus positions independent of deviant occurrence or learned expectations. **(E)** Response threshold. Higher values indicate a more conservative decision criterion, reflecting greater evidence accumulation before committing to a response. **(F)** Non-decision time. Higher values indicate that a longer pre-decisional delay, reflecting combined sensory encoding and motor execution, was necessary to account for the offset between predicted decision times and observed response times. *p < 0.05, **p < 0.01, ***p < 0.001, ns = not significant.

Here, we implemented an expanded version of the standard DDM that decomposes evidence accumulation into three separable components: (1) A sensory drift component capturing the quality of evidence provided by both standard (*v_s_*) and deviant (*v_d_*) stimuli, with larger deviant-to-standard drift differences (*v_s-_v_d_*) reflecting bottom-up deviant salience; (2) a deviant expectation weight *(wH_c_(t))* derived from the conditional probability of deviant occurrence at each stimulus position, capturing the contribution of learned top-down predictions to evidence accumulation; and (3) a time-dependent urgency signal *(U(t))* representing the contribution of temporal hazard to within trial responding (**Fig 4A**). These components were simultaneously fit to each animal’s response times and non-responses. Parameters were optimized for each animal that was subjected to both baseline and catch phases (n = 9 WT, 7 KO), allowing phase-dependent changes in decision parameters to be captured. Model validation confirmed adequate parameter recovery and robustness from initialization starting points (**Supplemental Fig 4**). This model thus provides a principled framework for dissociating the contributions of bottom-up sensory processing, top-down predictive expectations, and temporal hazard to the observed genotype differences in behavioral performance.

The DDM revealed a clear dissociation in the relative contributions of these components across genotypes. *Fmr1* KO animals showed a larger deviant-to-standard drift difference compared to WT controls (**Fig 4B**; Mann-Whitney U test: U = 9, *p = 0.016), reflecting a relative increase in evidence accumulation upon deviant presentation. Conversely, deviant expectation weighting was significantly reduced in *Fmr1* KO animals, with expectation weights approaching zero in nearly all subjects (**Fig 4C**; Mann-Whitney U test: U = 51, *p = 0.026), indicating that position-dependent changes in deviant probability had little effect on the decision process in *Fmr1* KO animals despite the stronger sensory signal. *Fmr1* KO subjects also showed greater temporal hazard weighting (**Fig 4D**; Mann-Whitney U test: U = 11, *p = 0.031) and reduced response threshold (**Fig 4E**; Mann-Whitney U test: U = 52, *p = 0.031) compared to WT animals. These results are consistent with a WT decision-making strategy in which responses are gated by conservative evidence accumulation and guided by learned predictive expectations. In contrast, *Fmr1* KO animals appeared to rely more strongly on within-trial response pressure and the immediate perceptual salience of the deviant, with little contribution from accumulated predictive expectations. Notably, non-decision time was comparable between genotypes (**Fig 4F**; Mann-Whitney U test: U = 40, p = 0.408), arguing against differences in sensory encoding or motor execution as drivers of the phenotype. Together, these results provide a formal decision-theoretic account of the behavioral findings that is consistent with a hypopriors phenotype in which *Fmr1* KO animals rely predominantly on sensory immediacy while WT animals integrate both sensory evidence and predictive expectations when deciding whether and when to respond.

## DISCUSSION

In this study, we leveraged a probabilistic oddball detection paradigm to examine predictive processing in an *Fmr1* KO rat model of FXS. We first established that this task provides a quantitative behavioral readout of sensory expectation. WT rats formed precise anticipatory representations of deviant occurrence that adaptively updated when task statistics changed, demonstrating that animals both form and flexibly maintain internal models of environmental contingencies. In contrast, *Fmr1* KO rats exhibited shallower position-dependent reaction time slopes and less precise false alarm distributions that remained relatively stable across task phases, suggesting deficits in both the formation and updating of predictive models. Drift diffusion modeling revealed that *Fmr1* KO animals showed near-absent expectation weighting but enhanced temporal hazard and deviant-driven evidence accumulation, consistent with a decision-making strategy driven by sensory immediacy rather than learned predictions. Together, these results establish a tractable experimental framework for studying predictive processing in rodent models and suggest that predictive processing deficits are a core feature of FXS.

The probabilistic nature of the task used in this study offers several advantages over classic oddball paradigms. First, it allows for the dissociation of deviant detection from deviant expectation by providing experimental control over the conditional probability of deviant occurrence in a manner that is independent of local stimulus statistics. Second, the probe phase provides a behavioral readout of deviant expectation in the absence of an actual deviant stimulus, enabling the quantification of internal model precision through the temporal distribution of false alarms on unreinforced probe trials. These probe trials are themselves higher-order sequence violations, similar to global deviants in a global-local oddball paradigm (Bekinschtein et al., 2009), providing a naturalistic way to examine responses to higher-order statistical violations. Third, our modeling approach demonstrates that this task can quantitatively constrain decision-making parameters related to response threshold, temporal hazard, and the relative weighting of immediate sensory evidence and learned expectations. Together, these features make the paradigm well-suited for evaluating theoretical accounts of predictive processing in preclinical rodent models..

One such theoretical prediction is that ASD would coincide with an overweighting of sensory information and/or an underweighting of the internal model (Lawson et al., 2014; Van de Cruys et al., 2014). Specifically, it has been suggested that core ASD phenotypes can be explained through “hypopriors”, or reduced weighting of prior expectations relative to incoming sensory evidence (Pellicano and Burr, 2012). Our findings in *Fmr1* KO rats are indeed consistent with theoretical accounts of altered precision-weighting in ASD. The less precise false alarm distributions and reduced position-dependence of reaction times observed in *Fmr1* KO rats are concordant with an imprecise prior that fails to optimally constrain behavior. Computational modeling further supported this interpretation, demonstrating that *Fmr1* KO rats overweight incoming sensory evidence and underweight internally generated expectations of deviant occurrence, resulting in behavioral performance that is dictated by sensory immediacy. Such aberrant precision in predictive coding is likely to contribute to sensory processing difficulties, intolerance of uncertainty, and behavioral inflexibility commonly observed in FXS and ASD.

While we believe our results are indicative of predictive processing deficits in *Fmr1* KO animals, several alternative explanations for the observed behavioral differences warrant consideration, including altered learning, motivation, attention, and/or motor responsivity. Our experimental design incorporates several features that help evaluate these possibilities. First, *Fmr1* KO rats acquired the task at comparable rates to WT littermates across training phases and maintained performance above criteria throughout the experiment. These findings argue against a gross impairment in learning of task rules. Importantly, comparable acquisition rates do not exclude more subtle genotype-dependent differences in learning processes that support the formation and updating of internal predictive models. The fact that *Fmr1* KO animals can learn to detect deviant stimuli but fail to anticipate deviant occurrence suggests that the ability to acquire task rules and the ability to use learned statistical regularities to guide anticipatory behavior may depend on partially dissociable processes. Second, *Fmr1* KO and WT subjects showed comparable trial initiation rates and attempts per trial, measures that provide implicit indices of self-motivation and task engagement. We did see a trend towards increased false alarm rates in *Fmr1* KO animals, which could reflect differences in attention or response inhibition that have been reported in this population (Schmitt et al., 2019). However, this modest shift in overall response tendency is unlikely to fully account for the observed pattern of position- and phase-dependent behavioral differences, as these stable trait differences would be expected to produce consistent, phase-independent effects on task performance. Finally, our DDM found no significant difference in non-decision time between genotypes, providing model-based evidence that alterations to pre-decisional sensory encoding and motor execution are unlikely to account for the observed behavioral differences. Together, these converging lines of evidence support the interpretation that the behavioral differences observed in *Fmr1* KO rats reflect a specific deficit in the formation and updating of internal predictive models, rather than a generalized performance impairment.

Position-dependent changes in reaction time and false alarm rate could also reflect temporal hazard or response urgency rather than learned expectations. However, several features of our data argue against a purely temporal account. First, a temporal hazard account would predict that false alarms should increase monotonically across all positions due to increasing urgency or motor preparation. In WT animals, false alarm rates on probe trials remained relatively constant across the first three stimuli (where no deviant can occur), ramped up at position 4 (where a deviant can first occur), peaked sharply at position 6 (the most expected deviant location) and then declined thereafter. This non-monotonic distribution is difficult to explain by hazard alone and is more consistent with a learned internal model of a specific expected deviant location. Second, WT animals adaptively modified their reaction times and response distributions when task statistics changed during catch phase, whereas these changes were less pronounced in *Fmr1* KO animals. Because the within-trial temporal structure remained unchanged across phases, this phase-dependent modulation suggests that behavior was influenced by learned task statistics rather than temporal hazard alone. Finally, our DDM explicitly parameterized temporal hazard and expectation weighting as separable components of evidence accumulation. *Fmr1* KO animals exhibited greater temporal hazard but reduced expectation weighting compared to WT animals. Thus, KO animals did not simply show a generalized reduction in temporal sensitivity; instead, the relative contributions of temporal hazard and learned expectations to behavior differed between genotypes.

The greater temporal hazard weighting in *Fmr1* KO rats is consistent with prior reports of hyperactivity and impulsivity in this model (D’Elia et al., 2022; Ntoulas et al., 2024), and may contribute to their higher overall false alarm rates. However, greater temporal hazard weighting alone cannot account for the specific pattern of position-dependent differences in reaction time and false alarm distributions. Indeed, greater temporal hazard weighting would be expected to produce larger position-dependent changes relative to WT animals — the opposite of what was observed here. Rather, the shallower reaction time slopes and broader false alarm distributions in *Fmr1* KO animals are better accounted for by their near-absent expectation weighting, consistent with animals that fail to translate learned predictions into anticipatory evidence accumulation. Together, these results suggest that predictive expectation and temporal hazard may represent two dissociable strategies for guiding behavior in this task, with *Fmr1* KO subjects using a strategy that favors temporal hazard and sensory salience over top-down expectations. Notably, genotypic differences in reaction time and false alarm distributions were substantially reduced during catch phase when deviant occurrence was no longer predictable, suggesting that WT animals may adopt a more temporally driven strategy when the predictive structure of the task is disrupted.

Our results also provide a novel lens for interpreting prior work characterizing sensory processing abnormalities in *Fmr1* KO rodents. We have previously shown that *Fmr1* KO rats exhibit faster reaction times in a sound detection paradigm that is associated with enhanced cortical gain (Auerbach et al., 2021; Gauthier et al., 2025). Interpreted through a predictive coding framework, enhanced neuronal gain could cause incoming stimuli to be treated as overly reliable, biasing internal models towards sensory immediacy at the expense of accumulated prior expectations. Faster reaction times may be reflective of this increased sensory immediacy, with *Fmr1* KO rats spending less time to integrate information and compare it to internal models. Consistent with this, our DDM revealed that *Fmr1* KO animals exhibited a lower decision threshold compared to WT animals and a greater difference in drift rates between deviant and standard stimuli, reflecting stronger relative evidence accumulation upon deviant presentation. This enhanced deviant salience may represent a decision-theoretic correlate of the increased cortical gain previously reported in this model. Importantly, the faster absolute reaction times in *Fmr1* KO animals are unlikely to account for the difference in position-dependent changes in reaction time observed in this study, as we found no significant correlation between absolute reaction time at position 4 and the magnitude of change across positions in either genotype, arguing against a floor effect. This interpretation is further supported by our DDM results, which revealed comparable non-decision times between genotypes, suggesting that the faster absolute reaction times in *Fmr1* KO animals reflect differences in the decision process itself rather than in motor execution. Together, these results suggest that increased sensory immediacy in *Fmr1* KO animals manifests as both faster responding and stronger bottom-up evidence accumulation at the expense of learned predictive expectations.

Our findings are also consistent with reports of impaired habituation to a repetitive stimulus in *Fmr1* KO mice and humans (Ethridge et al., 2016; Lovelace et al., 2016; He et al., 2017). Habituation is thought to reflect the compression of redundant sensory input into efficient internal models (Merchie and Gomot, 2023; Tsukano et al., 2026). Habituation deficits could thus be another outcome of impaired internal sensory model formation in FXS, leading to the continuous encoding of each stimulus with full fidelity, with this sensoryimmediacy coming at the expense of computational efficiency. Notably, the conservation of auditory habituation and evoked response phenotypes across FXS individuals and *Fmr1* KO rodents underscores the translational relevance of the current findings, suggesting that the current probabilistic oddball paradigm could provide a sensitive biomarker for sensory dysfunction in FXS individuals.

Sensory processing abnormalities in FXS are not restricted to the auditory domain either, with similar findings reported across somatosensory (He et al., 2017; Carreno-Munoz et al., 2018; Bhaskaran et al., 2023), visual (Goel et al., 2018; Pak et al., 2021), and olfactory (Kuruppath et al., 2023) modalities. Recent work has also demonstrated impaired sensory history integration in *Fmr1* KO mice during a tactile perceptual decision-making task (Semelidou et al., 2026), consistent with our finding of disproportionate weighting of immediate sensory evidence. The convergence of impaired predictive modeling with known sensory abnormalities in FXS suggests these phenomena may reflect a common underlying computational dysfunction rather than independent perceptual deficits.

Several limitations of the current study should be noted. First, our assessment was restricted to male subjects. While FXS is an X-linked disorder that occurs more frequently and with greater severity in males (Reiss and Hall, 2007), female FXS carriers can also present with a range of symptoms and severity due to mosaic X-inactivation (Bartholomay et al., 2019). It will therefore be important for future studies to determine if the differences in predictive coding observed here generalize to female *Fmr1* mutants. Additionally, while the current paradigm provides a sensitive behavioral readout of predictive processing, the neural mechanisms underlying these deficits remain to be characterized. Future work must address whether predictive deficits emerge from altered circuit properties like cortical hyperexcitability and degraded tuning (Rotschafer and Razak, 2013; Contractor et al., 2015; Goel et al., 2018; Antoine et al., 2019; Gauthier et al., 2025), or whether disrupted prediction itself drives these downstream changes. Resolving these mechanistic questions will be crucial for developing targeted interventions that address the root computational dysfunctions in FXS.

In summary, this study establishes a probabilistic oddball paradigm as a powerful tool for quantifying predictive processing in rodent models and provides behavioral and computational evidence that predictive processing is fundamentally disrupted in FXS. Beyond FXS, this paradigm offers a broadly applicable framework for interrogating the computational principles underlying perception across animal models of neuropsychiatric conditions, with clear translational parallels to oddball and mismatch negativity paradigms widely used in human clinical research. Sensory processing abnormalities are among the most prevalent features of ASD, observed not only in FXS but across other monogenic causes of ASD (Hudac et al., 2024), as well as in idiopathic autism (Kirby et al., 2022). Applying this paradigm across genetically diverse ASD models would allow for direct comparison of predictive processing profiles, testing whether sensory immediacy and hypopriorism represent conserved mechanisms for sensory dysfunction in ASD, or whether distinct etiologies produce qualitatively different disruptions to the balance between top-down and bottom-up processing. Beyond ASD, this paradigm may also be informative in models of schizophrenia, where some predictive coding accounts propose an opposite pattern of dysfunction — an over-reliance on top-down predictions at the expense of incoming sensory evidence (Sterzer et al., 2018; Qela et al., 2025). The quantitative behavioral and computational readouts afforded by this paradigm would provide a sensitive and tractable tool for testing this hypothesis in preclinical models, and more broadly for characterizing how the balance between top-down and bottom-up processing is disrupted across neuropsychiatric conditions with distinct etiologies.

## ACKNOWLEDGEMENTS

This work was supported by grants from the National Institutes of Health (R01HD111753, B.D.A.) and the Arnold and Mabel Beckman Foundation Graduate Fellowship (D.W.G.). We wish to acknowledge Dr. Xiaopeng Liu for his technical assistance on this project.

## Author contributions

Conceptualization: D.W.G., N.J., B.D.A.; Data Collection: D.W.G., N.J.; Analysis and Modeling: D.W.G., E.H., C.K.; Writing: D.W.G., B.D.A.

