## Supplemental Table 1 for "Impaired formation and updating of internal predictive models in a rat model of Fragile X Syndrome"

**Supplemental Table 1. Statistical analysis for all figure panels**

| **Figure** | **Outcome** | **Effect or comparison** | **Test** | **Statistic (df)** | **p** | **Effect size** |
| --- | --- | --- | --- | --- | --- | --- |
| 1B | Total training days | WT vs KO | Mann–Whitney | U = 64.500 | 1.000 | *r*_rb_ = −0.008 |
| 1C | Overall hit rate (%) | WT vs KO | Welch t | t(20.953) = 1.279 | 0.215 | g = 0.497 |
| 1D | Overall FA rate (%) | WT vs KO | Mann–Whitney | U = 31 | 0.060 | *r*_rb_ = −0.483 |
| 1E | Hit rate (%) | Genotype | Mixed ANOVA | F(1, 21) = 0.945 | 0.342 | *η*_p_^2^ = 0.043 |
| **1E** | **Hit rate (%)** | **Position** | **Mixed ANOVA** | **F(2, 42) = 95.853** | **<0.0001** | ***η*_p_^2^ = 0.820** |
| 1E | Hit rate (%) | Genotype × Position | Mixed ANOVA | F(2, 42) = 3.124 | 0.055 | *η*_p_^2^ = 0.129 |
| 1E | FA rate (%) | Genotype | Mixed ANOVA | F(1, 21) = 3.908 | 0.061 | *η*_p_^2^ = 0.157 |
| **1E** | **FA rate (%)** | **Position** | **Mixed ANOVA** | **F(2, 42) = 115.996** | **<0.0001** | ***η*_p_^2^ = 0.847** |
| 1E | FA rate (%) | Genotype × Position | Mixed ANOVA | F(2, 42) = 0.452 | 0.640 | *η*_p_^2^ = 0.021 |
| **1F** | **Normalized RT (ms)** | **Genotype** | **Mixed ANOVA** | **F(1, 21) = 6.644** | **0.012** | ***η*_p_^2^ = 0.249** |
| **1F** | **Normalized RT (ms)** | **Position** | **Mixed ANOVA** | **F(1, 21) = 64.320** | **<0.0001** | ***η*_p_^2^ = 0.754** |
| **1F** | **Normalized RT (ms)** | **Genotype × Position** | **Mixed ANOVA** | **F(1, 21) = 4.673** | **0.015** | ***η*_p_^2^ = 0.182** |
| **1G** | **Normalized RT slope (ms/position)ᵃ** | **WT vs KO** | **Welch t** | **t(17.509) = −3.119** | **0.006** | **g = −1.155** |
| **2B** | **Overall hit rate (%)** | **WT: baseline vs probe** | **Paired t** | **t(7) = −2.992** | **0.020** | ***d*_z_ = −1.058** |
| 2B | Overall hit rate (%) | KO: baseline vs probe | Paired t | t(6) = −0.411 | 0.695 | *d*_z_ = −0.155 |
| 2C | Overall FA rate (%) | WT: baseline vs probe | Wilcoxon | W = 18 | 1.000 | *r*_rb_ = 0.000 |
| 2C | Overall FA rate (%) | KO: baseline vs probe | Paired t | t(6) = −0.862 | 0.422 | *d*_z_ = −0.326 |
| **2E** | **Normalized RT (ms)** | **Genotype** | **Mixed ANOVA** | **F(1, 13) = 6.774** | **0.022** | ***η*_p_^2^ = 0.343** |
| **2E** | **Normalized RT (ms)** | **Position** | **Mixed ANOVA** | **F(2, 26) = 30.378** | **<0.001** | ***η*_p_^2^ = 0.700** |
| **2E** | **Normalized RT (ms)** | **Genotype × Position** | **Mixed ANOVA** | **F(2, 26) = 7.826** | **0.002** | ***η*_p_^2^ = 0.376** |
| **2F** | **Normalized RT slope (ms/position)ᵃ** | **Genotype** | **Mixed ANOVA** | **F(1, 13) = 11.938** | **0.004** | ***η*_p_^2^ = 0.479** |
| 2F | Normalized RT slope (ms/position)ᵃ | Phase | Mixed ANOVA | F(1, 13) = 0.204 | 0.659 | *η*_p_^2^ = 0.015 |
| 2F | Normalized RT slope (ms/position)ᵃ | Genotype × Phase | Mixed ANOVA | F(1, 13) = 0.660 | 0.431 | *η*_p_^2^ = 0.048 |
| **2H** | **FA time distribution** | **WT vs KO Probe** | **Cluster-permutation KS** | **D = 0.192** | **0.028** | **D = 0.192** |
| 3B | Overall hit rate (%) | Genotype | Mixed ANOVA | F(1, 14) = 0.844 | 0.374 | *η*_p_^2^ = 0.057 |
| 3B | Overall hit rate (%) | Phase | Mixed ANOVA | F(1, 14) = 0.557 | 0.468 | *η*_p_^2^ = 0.038 |
| 3B | Overall hit rate (%) | Genotype × Phase | Mixed ANOVA | F(1, 14) = 0.023 | 0.881 | *η*_p_^2^ = 0.002 |
| 3C | Overall FA rate (%) | Genotype | Mixed ANOVA | F(1, 14) = 2.366 | 0.146 | *η*_p_^2^ = 0.145 |
| 3C | Overall FA rate (%) | Phase | Mixed ANOVA | F(1, 14) = 0.509 | 0.487 | *η*_p_^2^ = 0.035 |
| **3C** | **Overall FA rate (%)** | **Genotype × Phase** | **Mixed ANOVA** | **F(1, 14) = 6.315** | **0.018** | ***η*_p_^2^ = 0.311** |
| **3D** | **Normalized RT (ms), WT** | **Phase** | **RM ANOVA** | **F(1, 8) = 34.806** | **<0.001** | ***η*_p_^2^ = 0.813** |
| **3D** | **Normalized RT (ms), WT** | **Position** | **RM ANOVA (GG)** | **F(1.138, 9.104) = 8.462** | **0.015** | ***η*_p_^2^ = 0.514** |
| 3D | Normalized RT (ms), WT | Phase × Position | RM ANOVA (GG) | F(1.156, 9.252) = 31.874 | <0.001 | *η*_p_^2^ = 0.799 |
| 3E | Normalized RT (ms), KO | Phase | RM ANOVA | F(1, 6) = 3.812 | 0.066 | *η*_p_^2^ = 0.175 |
| **3E** | **Normalized RT (ms), KO** | **Position** | **RM ANOVA (GG)** | **F(1.326, 7.957) = 9.116** | **0.013** | ***η*_p_^2^ = 0.603** |
| 3E | Normalized RT (ms), KO | Phase × Position | RM ANOVA (GG) | F(1.482, 8.893) = 1.120 | 0.331 | *η*_p_^2^ = 0.157 |
| 3F | Normalized RT slope (ms/position)ᵃ | Genotype | Mixed ANOVA | F(1, 14) = 0.237 | 0.634 | *η*_p_^2^ = 0.017 |
| **3F** | **Normalized RT slope (ms/position)ᵃ** | **Phase** | **Mixed ANOVA** | **F(1, 14) = 15.765** | **0.001** | ***η*_p_^2^ = 0.530** |
| **3F** | **Normalized RT slope (ms/position)ᵃ** | **Genotype × Phase** | **Mixed ANOVA** | **F(1, 14) = 6.873** | **0.020** | ***η*_p_^2^ = 0.329** |
| 3G | FA time distribution | WT Probe vs Catch | Cluster-permutation KS | D = 0.266 | 0.032 | D = 0.266 |
| 3H | FA time distribution | KO Probe vs Catch | Cluster-permutation KS | D = 0.119 | 0.124 | D = 0.119 |
| **3I** | **Percentage of FAs in position-6 window** | **Genotype** | **Two-way ANOVA** | **F(1,27) = 10.93** | **0.003** | **0.288** |
| **3I** | **Percentage of FAs in position-6 window** | **Phase** | **Two-way ANOVA** | **F(1,27) = 8.79** | **0.006** | **0.247** |
| 3I | Percentage of FAs in position-6 window | Genotype × Phase | Two-way ANOVA | F(1,27) = 2.678 | 0.114 | 0.090 |
| **4B** | **Deviant salience (*v*_d_ − *v*_s_)** | **WT vs KO** | **Mann–Whitney (P)** | **U = 9** | **0.016** | ***r*_rb_ = −0.714** |
| **4C** | **Expectation weighting (w)** | **WT vs KO** | **Mann–Whitney (P)** | **U = 51** | **0.026** | ***r*_rb_ = 0.619** |
| **4D** | **Temporal response pressure slope (u)** | **WT vs KO** | **Mann–Whitney (P)** | **U = 11** | **0.031** | ***r*_rb_ = −0.651** |
| **4E** | **Response threshold (a)** | **WT vs KO** | **Mann–Whitney (P)** | **U = 52** | **0.031** | ***r*_rb_ = 0.651** |
| 4F | Nondecision time (s) | WT vs KO | Mann–Whitney (P) | U = 40 | 0.408 | *r*_rb_ = 0.270 |
| S1A | Detection attempts/trial | WT vs KO | Mann–Whitney (BH) | U = 61 | 0.876 | *r*_rb_ = −0.062 |
| S1A | Detection training days | WT vs KO | Mann–Whitney (BH) | U = 80 | 0.662 | *r*_rb_ = 0.231 |
| S1B | Discrimination attempts/trial | WT vs KO | Mann–Whitney (BH) | U = 46 | 0.805 | *r*_rb_ = −0.292 |
| S1B | Discrimination training days | WT vs KO | Welch t (BH) | t(12.823) = −0.865 | 0.805 | g = −0.379 |
| S1C | Oddball attempts/trial | WT vs KO | Welch t (BH) | t(20.031) = 0.158 | 0.876 | g = 0.063 |
| S1C | Oddball training days | WT vs KO | Mann–Whitney (BH) | U = 57.500 | 0.876 | *r*_rb_ = −0.115 |
| **S3A** | **Raw RT (ms)** | **Genotype** | **Mixed ANOVA** | **F(1, 21) = 5.739** | **0.026** | ***η*_p_^2^ = 0.215** |
| **S3A** | **Raw RT (ms)** | **Position** | **Mixed ANOVA (GG)** | **F(1.085, 22.795) = 60.157** | **<0.001** | ***η*_p_^2^ = 0.741** |
| **S3A** | **Raw RT (ms)** | **Genotype × Position** | **Mixed ANOVA (GG)** | **F(1.085, 22.795) = 5.671** | **0.024** | ***η*_p_^2^ = 0.213** |
| S3B | Position-4 RT vs position-6 speeding (ms)ᵇ | WT | Pearson r | r(8) = 0.117 | 0.747 | r = 0.117 |
| S3B | Position-4 RT vs position-6 speeding (ms)ᵇ | KO | Pearson r | r(11) = 0.038 | 0.903 | r = 0.038 |
| S4A | **Standard drift (*v*_s_)** | **WT vs KO** | **Mann-Whitney (P)** | **U = 52** | **0.031** | **r_rb_ = 0.651** |
| S4A | Deviant drift (*v*_d_) | WT vs KO | Mann-Whitney (P) | U = 41 | 0.351 | *_r_*_rb_  = 0.302 |
