## Supplemental Figures for "Impaired formation and updating of internal predictive models in a rat model of Fragile X Syndrome"

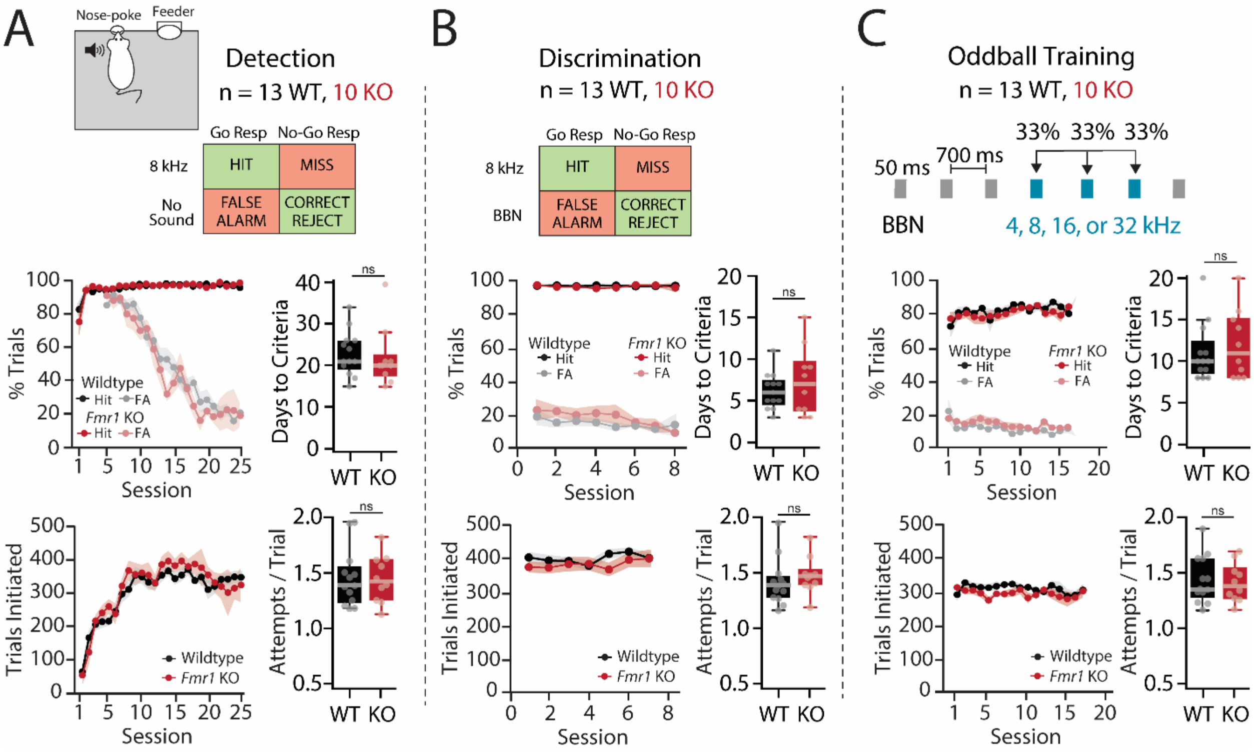


**Supplemental Figure 1.** Behavioral training across **(A)** Detection; **(B)** Discrimination; and **(C)** Oddball task phases. WT (n = 13) and *Fmr1* KO (n = 10) animals exhibited similar hit rates, false alarm (FA) rates, and trials initiated per session for across all three phases of training. There was no significant difference in days to criteria for any phase (Detection: Mann-Whitney, U = 80, p = 0.662; Discrimination: Welch's t-test, t(12.823), p = 0.805; Oddball: Mann-Whitney, U = 57.5, p = 0.876). Additionally, there was no difference in attempts/trial for any training phase (Detection: Mann-Whitney, U = 61, p = 0.876; Discrimination: Mann-Whitney, U = 46, p = 0.805; Oddball: Welch's t-test, t(20.031), p = 0.876), indicating similar levels of task engagement and motivation.

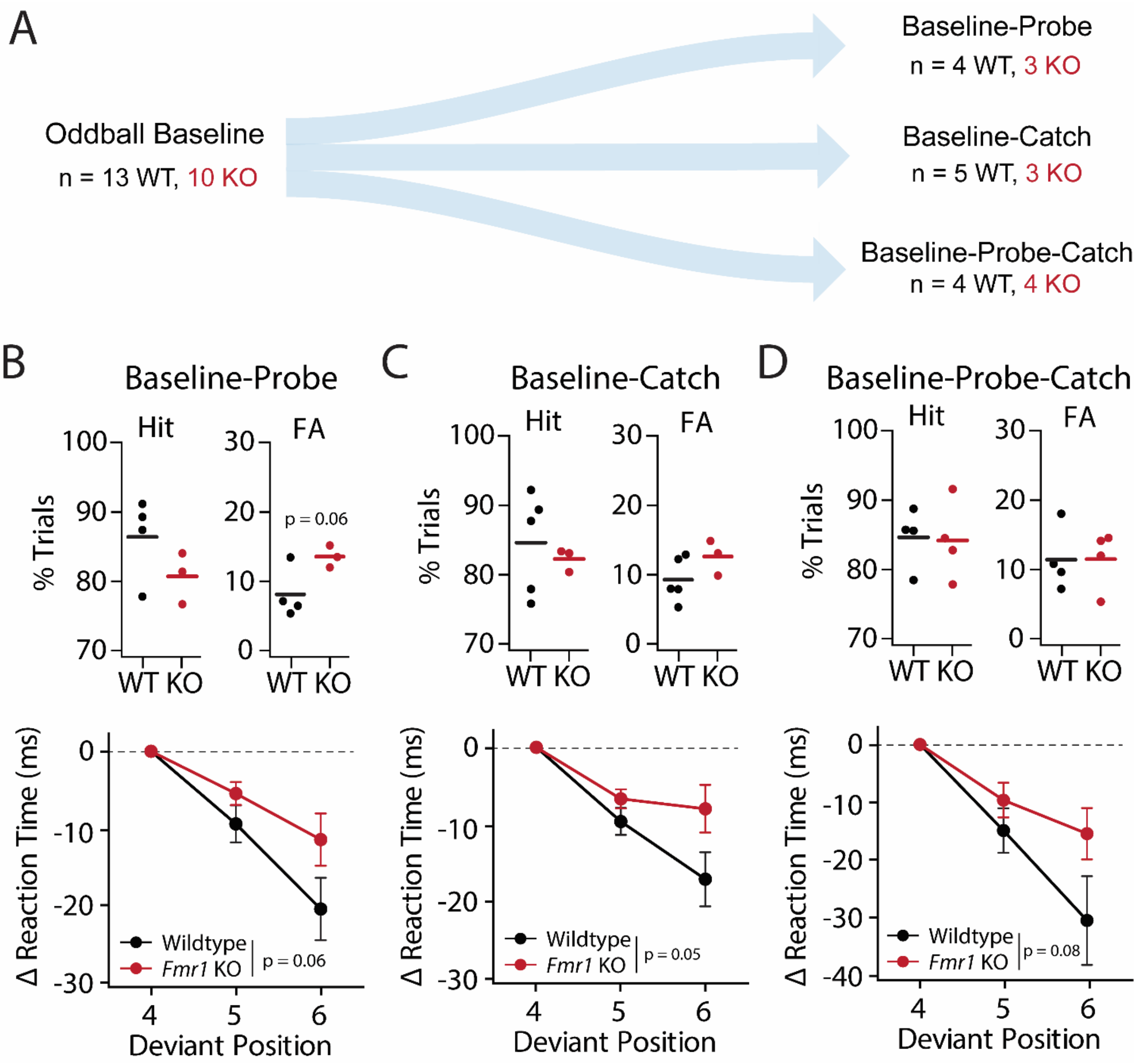


**Supplemental Figure 2.** Experimental design. **(A)** A total of 13 WT and 10 *Fmr1* KO rats were tested on the baseline phase of the task across three cohorts. Cohort 1 was subjected to probe phase (but not catch phase) following baseline testing, while cohort 2 was subjected to catch phase (but not probe phase). Cohort 3 underwent all three phases (baseline, probe, catch). This resulted in an overlapping yet unique set of animals for each phase. **(B-D)** Hit rates, false alarm (FA) rates, and normalized reaction time x position curves for each cohort during baseline phase of testing demonstrated qualitatively similar behavioral performance across cohorts.

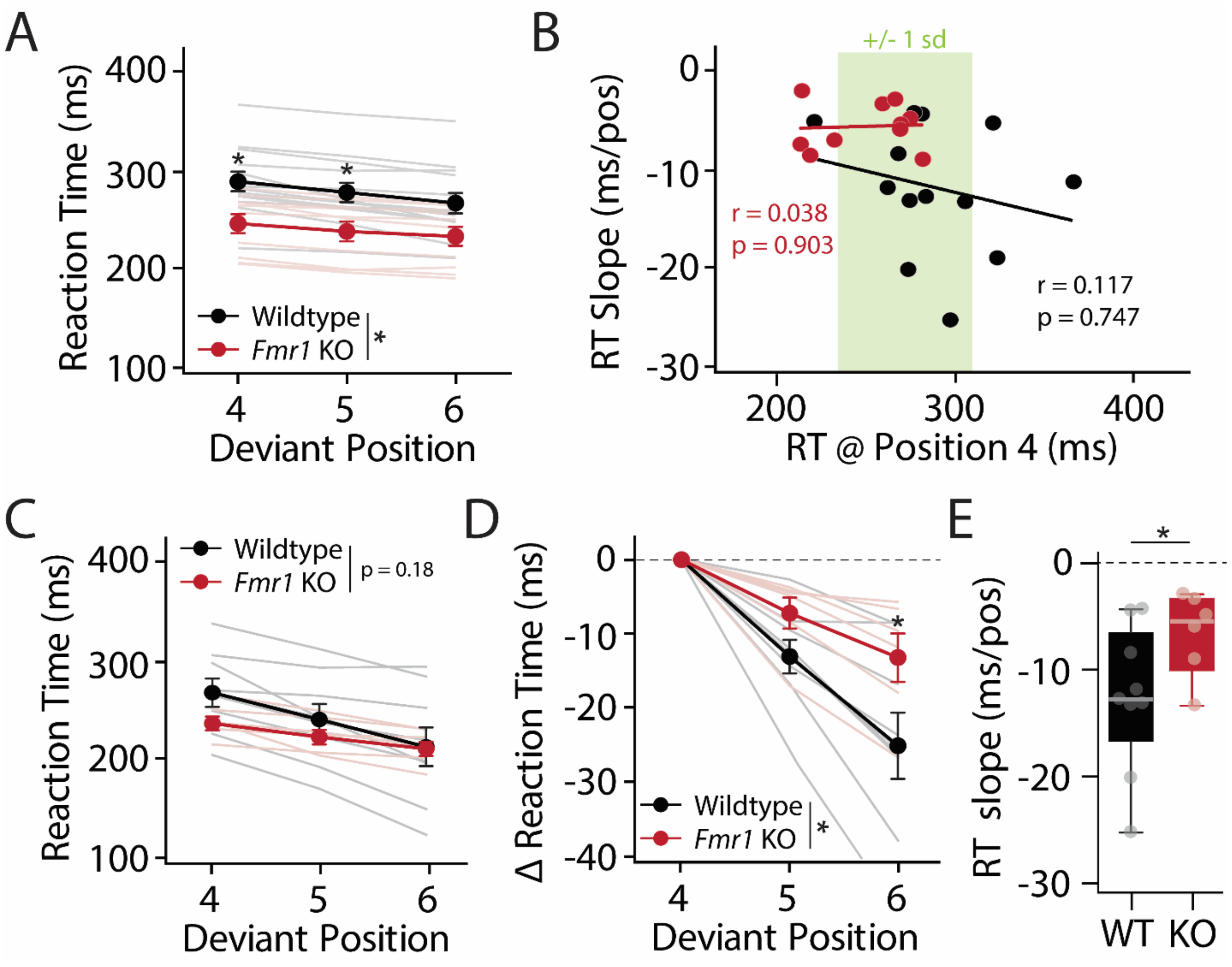


**Supplemental Figure 3.** Relationship between absolute reaction time (RT) and position-dependent changes in RT across genotypes. **(A)** Absolute RT as a function of deviant position in wildtype (n = 13) and Fmr1 KO (n = 10) rats during baseline phase of testing. *Fmr1* KO rats exhibited significantly faster RTs with most pronounced differences for deviants occurring at position 4 and 5 (mixed ANOVA: genotype, $F(1,21) = 5.739$, *$p = 0.026$). **(B)** No significant correlation was observed between absolute RT in response to position 4 deviants and the magnitude of RT change across positions (RT slope) in either genotype (WT: black, KO: red), arguing against a floor effect. If reduced RT slope in *Fmr1* KO animals were a consequence of already-fast reaction times, it would be expected that subjects with the fastest absolute RTs would show the least position-dependent change. **(C)** Absolute and **(D)** position-normalized reaction time curves for subset of animals (WT: n = 9, KO: n = 6) whose position 4 reaction time was within +/- 1 standard deviation of mean position 4 reaction time across genotypes (green shading in B). While there was no significant difference in absolute RT curves for this subgroup (mixed ANOVA: genotype,

$F(1,13) = 1.866$, $p = 0.180$), position normalized RT curves (mixed ANOVA: genotype, $F(1,13) = 6.542$, *$p = 0.015$) and **(E)** RT slopes (Welch's t-test, $t(13) = 1.951$, * $p = 0.037$) were still significantly different, indicating that deviant position differences in reaction time is not dependent on absolute RT.

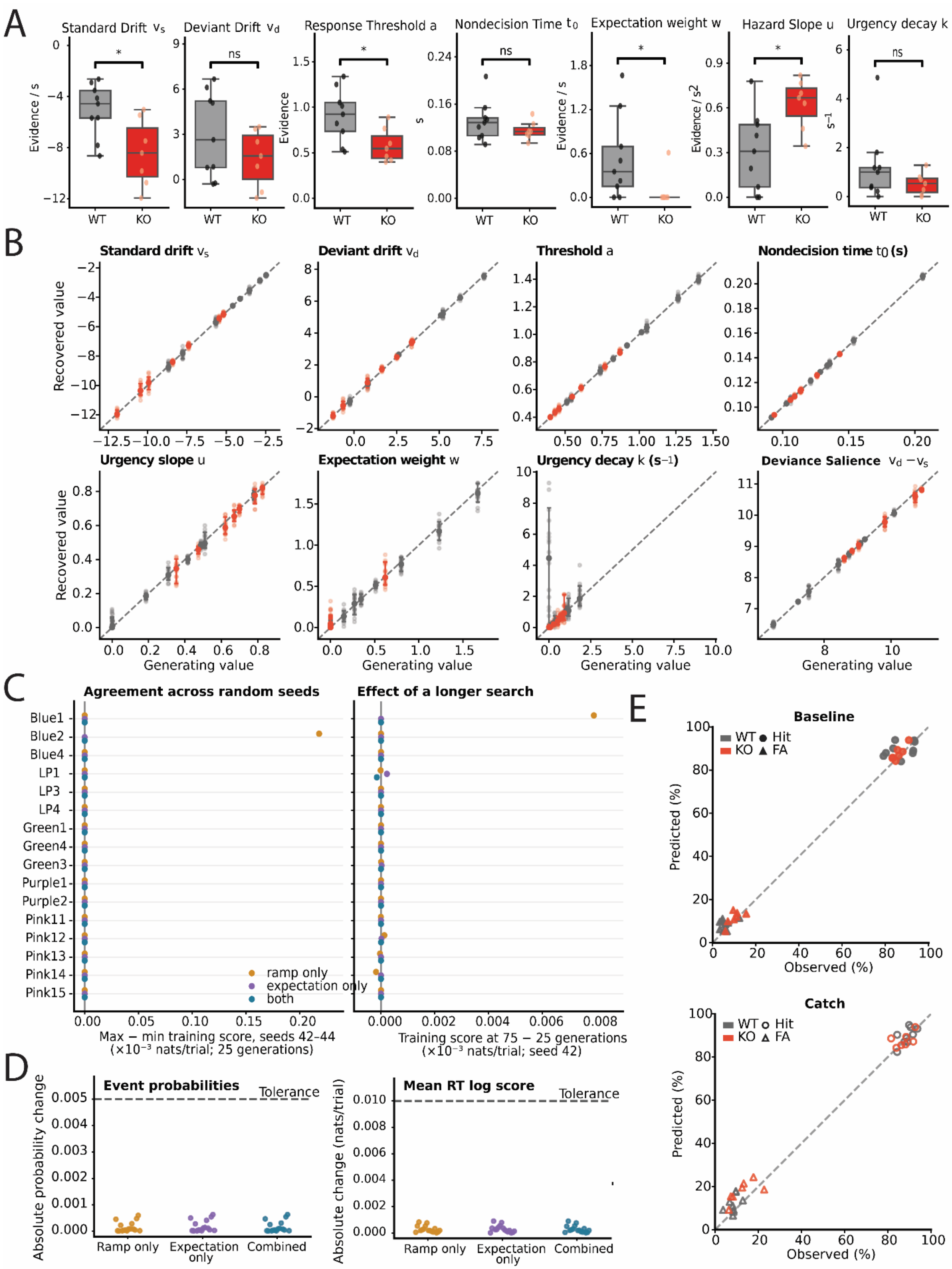

A
Standard Drift $v_s$
Deviant Drift $v_d$
Response Threshold a
Nondecision Time $t_0$
Expectation weight w
Hazard Slope u
Urgency decay k
Evidence / s
Evidence
s
Evidence / s
Evidence / s2
s−1
WT
KO
*
ns
B
Standard drift $v_s$
Deviant drift $v_d$
Threshold a
Nondecision time $t_0$ (s)
Urgency slope u
Expectation weight w
Urgency decay k (s−1)
Deviance Salience $v_d - v_s$
Recovered value
Generating value
C
Agreement across random seeds
Effect of a longer search
Blue1
Blue2
Blue4
LP1
LP3
LP4
Green1
Green4
Green3
Purple1
Purple2
Pink11
Pink12
Pink13
Pink14
Pink15
ramp only
expectation only
both
Max − min training score, seeds 42–44
(×10−3 nats/trial; 25 generations)
Training score at 75 − 25 generations
(×10−3 nats/trial; seed 42)
D
Event probabilities
Mean RT log score
Tolerance
Absolute probability change
Absolute change (nats/trial)
Ramp only
Expectation only
Combined
E
Baseline
Catch
WT
KO
Hit
FA
Predicted (%)
Observed (%)

**Supplemental Figure 4.** Drift Diffusion Model Validation. **(A)** All fitted parameters for WT (n = 9) and *Fmr1* KO (n = 7) subjects. **(B)** Model parameter recovery. 20 simulated datasets per animal using its fitted parameters and training-condition counts, then freely refitted all seven parameters from randomized starting populations (320 datasets; 9 WT, 7 KO). Most parameters were recovered close to the identity line, indicating agreement between generating and estimated values. Decay was the main exception: A single WT rat generating decay of zero yielded a median estimate of 4.46, consistent with weak information about decay when urgency is near zero. Faint dots: individual refits. Large dots: animal medians. Whiskers: empirical 10-90$^{th}$ percentiles. **(C)** Initialization (16 animals x 3 model structures; global searches followed by local refinement). Different random seeds and longer optimization searches produced similar training likelihoods. Across the 48 animal/model combinations, the largest between-seed score difference was approximately 0.000218 nats per trial, below the predefined 0.001 threshold. Because searches included shared informed starting vectors, this demonstrates stability under the implemented warm-start procedure rather than fully independent initialization or uniquely determined parameters. **(D)** Resolution. Increasing the evidence grid from 80 to 160 cells and reducing the time step from 2.5 to 1.25 ms produced minimal changes in predictions. All 48 checks passed, with maximum changes of 0.000633 in event probability and 0.000903 nats per trial in mean response-time log score. Predictions were therefore stable across the numerical resolutions tested. Each dot is one animal/model combination. **(E)** Predicted versus observed hit rates and false alarm rates in WT (black) and Fmr1 KO (red) animal during baseline (top) and catch (bottom) phases using held-out evaluation sessions. Each point represents one animal; dashed line indicates perfect correspondence.
